# Combining molecular and cell painting image data for mechanism of action prediction

**DOI:** 10.1101/2022.10.04.510834

**Authors:** Guangyan Tian, Philip J Harrison, Akshai P Sreenivasan, Jordi Carreras Puigvert, Ola Spjuth

**Affiliations:** Department of Pharmaceutical Biosciences, Uppsala University; Department of Medical Sciences, Uppsala University

## Abstract

The mechanism of action (MoA) of a compound describes the biological interaction through which it produces a pharmacological effect. Multiple data sources can be used for the purpose of predicting MoA, including compound structural information, and various assays, such as those based on cell morphology, transcriptomics and metabolomics. In the present study we explored the benefits and potential additive/synergistic effects of combining structural information, in the form of Morgan fingerprints, and morphological information, in the form of five-channel Cell Painting image data. For a set of 10 well represented MoA classes, we compared the performance of deep learning models trained on the two datasets separately versus a model trained on both datasets simultaneously. On a held-out test set we obtained a macro-averaged F1 score of 0.58 when training on only the structural data, 0.81 when training on only the image data, and 0.92 when training on both together. Thus indicating clear additive/synergistic effects and highlighting the benefit of integrating multiple data sources for MoA prediction.

## Introduction

Mechanism of action (MoA) refers to the biological interaction through which a potentially therapeutic small-molecule compound produces a pharmacological effect, such as the specific proteins that the compound targets and the pathways that it modulates. Uncovering the MoA of a compound, although a significant challenge in chemical biology [1], provides extremely useful information for lead compounds prior to clinical trials and for identifying possible toxicity or side-effects [2].

A variety of different data sources can be used to capture information on a compound’s MoA, including structural information from the compound, gene expression from transcriptomics data, protein information from proteomics data, and metabolic enzyme activity from metabolomics data [2]. Recently, cell morphology data from high-content imaging has proven useful for this task [3]. A significant benefit of microscopy based image assays is that they can be scaled to high-throughput much more easily and less expensively than transcriptomics and metabolomics based assays [4]. Cell imaging also provides information at the single-cell resolution as opposed to condensing the output down to measures of population averages [5]. In terms of throughput and efficiency the L1000 [6] gene expression assay is perhaps currently the only feasible alternative to image-based assays [7] for large scale data generation to sustain predictive modeling.

Microscopy imaging can be used to capture the changes in cell morphology that arise when a cell culture is treated with a chemical compound [2]. The Cell Painting assay uses fluorescent dyes to paint the cells in multiwell plates as “richly as possible” to illuminate morphological changes in eight broadly relevant organelles and cellular sub-compartments (nuclei, mitochondria, cytoskeleton, Golgi apparatus, plasma membrane, cytoplasmic RNA, nucleoli and endoplasmic reticulum) using six fluorescent dyes imaged in five channels [7].

A comparative study for library enrichment reported better predictive power for High-throughput screening performance using Cell Painting as opposed to L1000 gene expression profiling [8]. Whereas, for predicting MoA, Way et al. [9] found that L1000 outperformed Cell Painting, but that there was complementarity, i.e. some MoAs were better predicted by one of the assays compared to the other. A related study by Lapins and Spjuth [10] compared Cell Painting, L1000 and chemical structure based predictors, and found MoA classes that were predicted better by each of the three predictors relative to the other two, supporting the idea of a likely benefit through combining these different data sources. Another study predicting MoA [11], based on data from the ExCAPE database, which compared models built using image based features to those built using chemical structure descriptors, provided further support for the complementarity of these two data types, whereby the models performed somewhat differently at an individual class level.

Besides comparing models built using different types of data it is also possible to combine the datasets and analyze them simultaneously to search for additive or synergistic effects. For predicting cytotoxicity and proliferation Seal et al. [12] compared Random Forest models using Cell Painting image based features, molecular fingerprints, and combining both data sources. They found that the models based on image features outperformed those based on molecular fingerprints, but the combined models performed best in ten out of twelve cases. Another study predicting the bio-activity of approximately 16,000 compounds [13], found that models based on features derived from Cell Painting images outperformed those based on chemical structure profiles from graph convolutional networks (GCNs, [14]), but that the fusion of the two datasets gave a gain in performance.

Most traditional image analysis pipelines, including those mentioned above, first extract morphological features from the fluorescence stained images, including measures of size, shape, intensity and texture from the labeled cellular compartments, most often using the CellProfiler [15] software package, and subsequently apply machine learning methods to the extracted features for the predictive task at hand [16]. These methods require an accurate segmentation algorithm to identify the cellular compartments prior to feature extraction. However, when convolutional neural networks (CNNs) are used on the raw images, features are extracted in an automatic data-driven fashion, circumventing the need for cell segmentation and potentially providing better predictive performance [3,17]. For instance, Hofmarcher et al. [18] found that CNNs trained on Cell Painting image data, for predicting activity labels for over 10,000 compounds, performed significantly better than fully connected neural networks trained on pre-computed image features. The flexibility of the architectural choices for neural networks also provides a simple means of combining multiple data sources into the same modeling framework [19].

In the work presented in this manuscript we first compared a variety of traditional machine learning and deep learning models for the prediction of MoA based on chemical structure data for up to 20 MoA classes. Subsequently, based on a set of 10 MoA classes, we compared the performance of the best deep learning model at the compound structural level to a state-of-the-art CNN trained on Cell Painting image data for the same set of compounds. We selected the best deep learning based compound structure model so that we could finally train a joint model for the MoA prediction based on utilizing both structural and image data as input. Example Cell Painting images for the 10 MoA classes can be seen in Figure 1. To the best of our knowledge our work represents the first combination of five channel Cell Painting image data and molecular fingerprint data trained in an end-to-end fashion to predict MoA, wherein the raw images, as opposed to features derived from the images, were used as input to the models.

**Figure 1.**
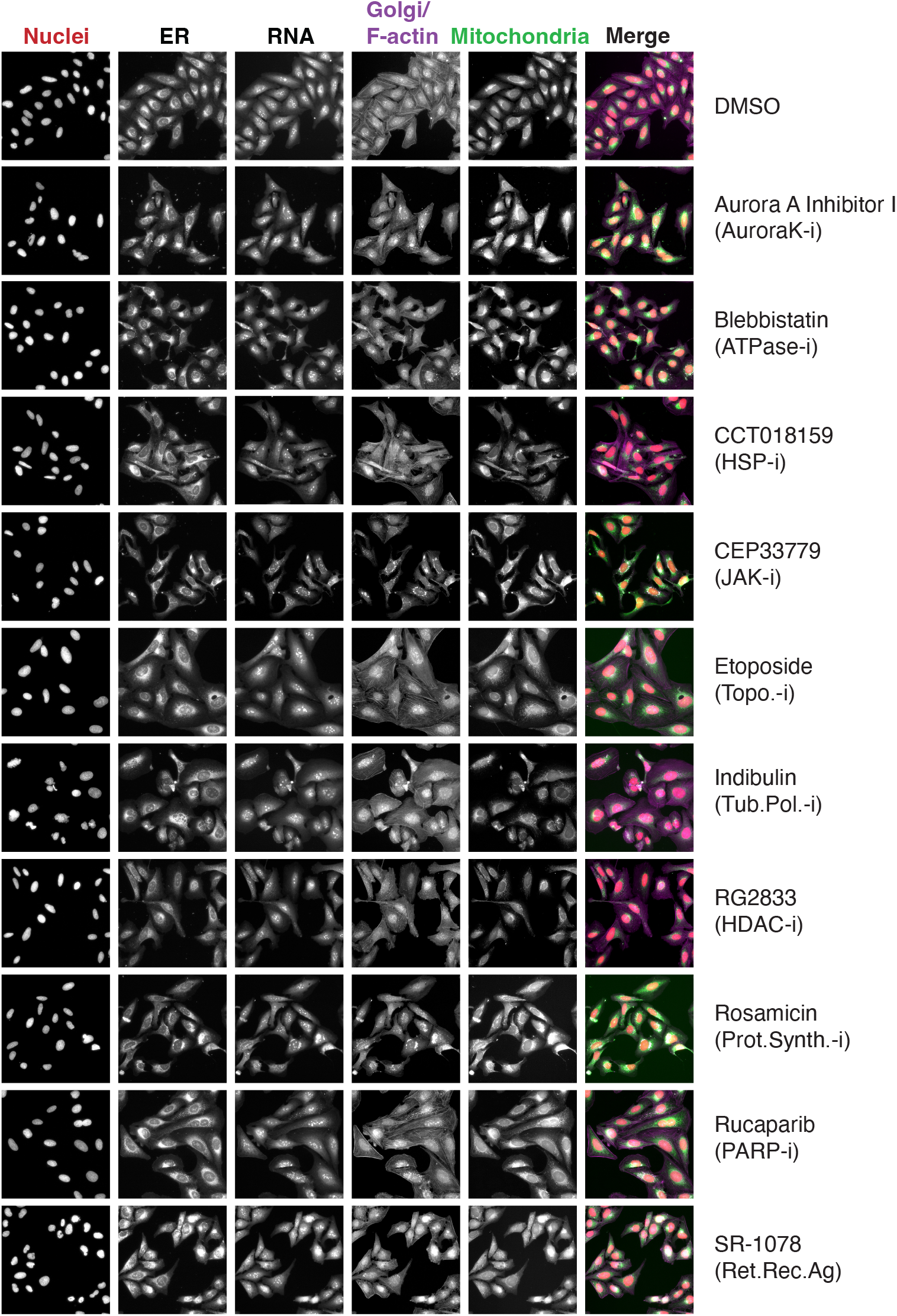
Example Cell Painting images for the 10 MoA classes and the DMSO data used for standardization. The row titles give the compound names for the selected images with the MoA abbreviation in parenthesis, where i stands for inhibitor and Ag for agonist.

## Materials and Methods

### Data

#### Molecular data

Molecular data (Corsello et al. [20]), in the form of SMILES strings collected and processed by the Broad Institute, was used in this study. The cleansed dataset contains approximately 5,500 compounds covering 1,300 MoA classes, but most MoAs have very few compounds associated with them. The number of compounds that each MoA has is shown in Figure 2. As our models should perform well at the compound level, namely to predict the MoA for unseen compounds, we used a subset of the data, the top 20 MoAs (i.e. the 20 MoAs having the most compounds associated with them).

**Figure 2.**
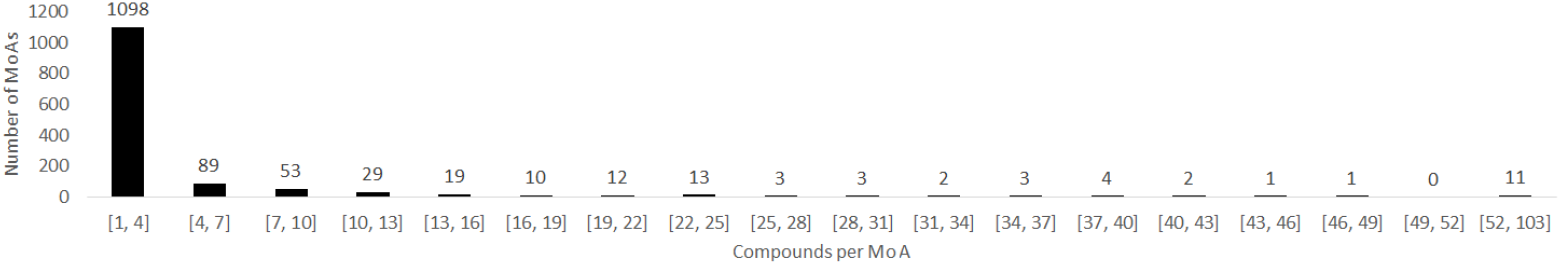
Histogram representing the compound counts per MoA for different binning intervals. Note that the range of the interval for the final bin is larger than the others.

#### Image data

The 5-channel Cell Painting image data was produced by the Pharmaceutical Bioinformatics Research Group at Uppsala University. We selected image data from 10 well-represented MoAs (MoAs that we presumed would be reasonably distinguishable and that had a sufficient number of compounds associated with them). The ten MoAs were Aurora kinase inhibitor (number of compounds, n = 20), tubulin polymerization inhibitor (n = 20), JAK inhibitor (n = 21), protein synthesis inhibitor (n = 23), HDAC inhibitor (n = 33), topoisomerase inhibitor (n = 32), PARP inhibitor (n = 21), ATPase inhibitor (n = 18), retinoid receptor agonist (n = 19), and HSP inhibitor (n = 24). In total we had 12582 images for 231 compounds. The compounds were administered to U2OS cells in 384 well plates at a dose of 10 micro-molar. Images at a resolution of 2160 × 2160 pixels were taken across 9 sites in each well and each compound was replicated 6 times. The compounds were distributed across 18 plates using PLAID (Plate Layouts using Artificial Intelligence Design, [21]).

#### Data preprocessing for models based on molecular data

We explored usin multi-layer perceptrons (MLPs), graph convolutional networks (GCNs), convolutional neural networks (CNNs), long short-term memory networks (LSTMs, with and without data augmentation), and traditional machine learning algorithms (those operating on tabular data) to predict MoA, and we pre-processed the data for each model. See the Modeling section below for further details on the models explored.

For the MLP and traditional machine learning algorithms, we used Morgan Fingerprints as input. Since SMILES strings are sequential they cannot be processed directly by these models. We used the RDKit package [22] to generate the Morgan fingerprints (binary vectors, 2048 bits) [23]. For the GCN, which requires the adjacency matrix and the node matrix as input, we applied Spektral [24] and NetworkX packages [25] to convert the SMILES strings into graphs. For the CNN we generated the feature matrix for each SMILES string based on the approach of Hirohara et al. [26]. Initially, we selected 42 chemical features to prepare the feature matrix of each SMILES string based on the selected chemical features. Secondly, due to the blank parts of the feature matrix resulting from inconsistent lengths of the SMILES, we applied zero padding to maintain uniform dimensions of the feature matrix. For the recurrent neural network, the LSTM, we utilized SMILES pair encoding [27] to tokenize the chemical structure data, so we obtained a series of numbers (tokens) representing the SMILES. Similarly to the CNN case, we also used zero padding to ensure that the lengths of all tokens were identical.

#### Data preprocessing for models based on image data

The 5 channels in the Cell Painting image data were standardized to remove plate-level effects based on the mean and standard deviation of the pixel intensities in the control DMSO wells in each plate. The images were resized from their original dimension down to 256 × 256 pixels. A quality control run on the data to detect saturation and blur in the images found no saturation issues (such as fibers across the field of view) but did detect some blurred images. However, given that a common data augmentation strategy for deep learning models is to purposefully blur the images we decided not to remove these blurred images from the dataset.

#### Data augmentation and data splitting

Data augmentation generates additional data based on the existing data and improves the generalizability of models. For the image based models we used flipping, 90 degree rotations and shift scale rotations to augment the data. However, for the compound structure based models data augmentation was only possible for the LSTM. As the LSTM is a sequence-based model that requires tokens as input, data augmentation is feasible as slightly different tokens can be produced by randomized SMILES [28].

For splitting the data at the compound-level into training, validation and test sets we used stratification based on the proportion of compounds for each MoA. We split out 10% of the data for the final held-out test set. Stratified splitting for the remaining data was performed nine times (nine shuffles) for the SMILES data in the initial comparison and five times (five shuffles) for the image data and the corresponding SMILES subset. In each case 80% of the data was used for training and 10% for validation.

### Modeling

#### Compound structure based models

We explored the following deep learning models for the prediction of MoA using chemical structure data: MLP, GCN, CNN, and LSTM with and without data augmentation. For the deep learning models we determined the optimal architectures and parameters through model exploration and parameter tuning on the validation sets. The MLP is a basic artificial neural network [29] that includes fully connected input, hidden, and output layers. Our MLP model contained one input layer, one hidden layer with dropout (p = 0.85), and one final prediction layer. GCNs are a subset of GNNs [30] that can process non-Euclidean data, such as graphs with nodes and edges [14]. Our GCN model included input layers for the adjacency matrix and the node matrix followed by three convolution layers with dropout (p = 0.5), one global attention pooling layer and one final prediction layer. Our CNN model contained one convolution layer, one max pooling layer with dropout (p = 0.8), one flattening layer with dropout (p = 0.8), and one final prediction layer. Our LSTM model included an embedding layer, a bidirectional LSTM layer, a dropout layer (p = 0.96) and a final prediction layer. For the LSTM with data augmentation, we adjusted the degree of augmentation to ensure that each MoA had approximately 1000 SMILES in the augmented training set.

We used the Adam optimizer [31], sparse categorical cross-entropy as the loss function, and validation loss as the metric for early stopping. To accommodate for imbalance of classes we applied class weighting in the loss functions to train the models.

We also explored machine learning algorithms that operate on tabular data (in contrast to the deep neural networks described above). The more traditional machine learning models have shown competitive performance with deep learning models when dataset sizes are relatively small [32]. For instance, Jiang et al. [33] showed that four descriptor-based models outperformed four graph-based models on several benchmark datasets. We examined five individual machine learning algorithms and four ensemble algorithms. The individual algorithms included random forests [34], light gradient boosting machines [35], cat boost [36], k-nearest neighbors classifiers [37], and logistic regression [38]. The ensemble algorithms included bagging [39], stacking [40], voting [41], and adaboost [42].

#### Cell morphology based model

We applied the state-of-the-art CNN model EfficientNet [43] to predict MoA based on the 5-channel Cell Painting image data. EfficientNet applies a compound scaling method to adjust width, depth, and resolution simultaneously, achieving competitive performance in image-based tasks with less training time and fewer parameters. We adopted the EfficientNetB1 architecture and used the AdamW optimizer [44] with weigthed sparse categorical cross entropy as the loss function.

#### Global model

For our global model trained on the data for the 10 selected MoA classes, we integrated the MLP (our best performing deep learning model based on the compound structure data) and EfficientNet (for the image data). The models were first trained separately and then combined and their weights finetuned. The architecture of our global model is shown in Figure 3.

**Figure 3.**
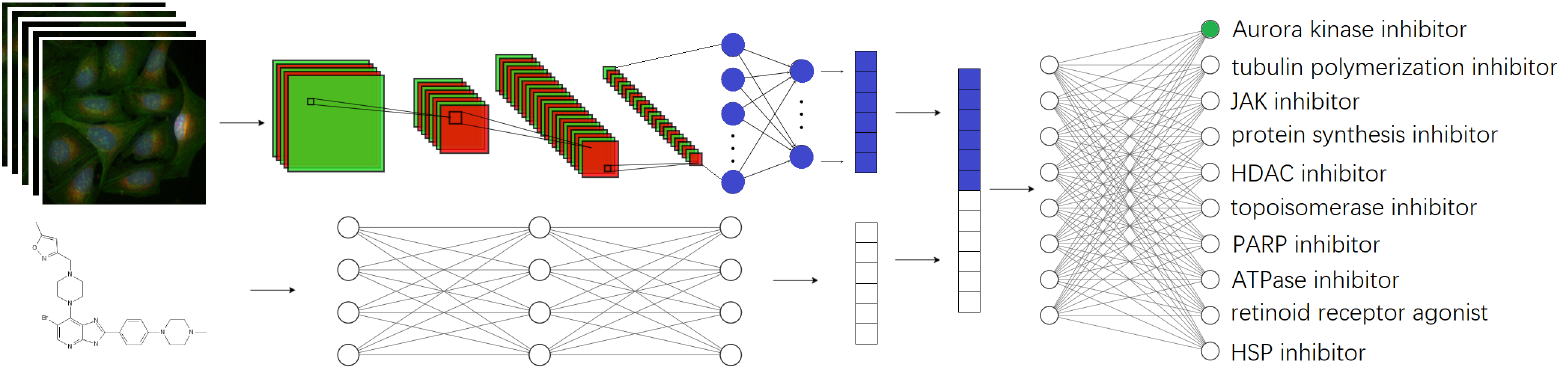
The architecture of the global model with two input paths, one for the Cell Painting image data and one for the chemical structure data.

## Results

Summaries of the performance of the traditional and deep learning models for the compound structure based models for the 20 MoA subsets are shown in Figure 4. These figures present the average F1 scores across the nine shuffles of the training and validation data as well as the results of randomisation tests performed to assess the level of significance in the performance differences. We applied Bonferroni corrections [45] to the p-values to account for the fact that we were performing several comparisons. The performances of the traditional machine learning algorithms were all quite comparable, however there were larger differences for the deep learning models compared. The best performing deep learning model was the MLP and the worst was the CNN; we note that the MLP performed on par with the best traditional machine learning models.

**Figure 4.**
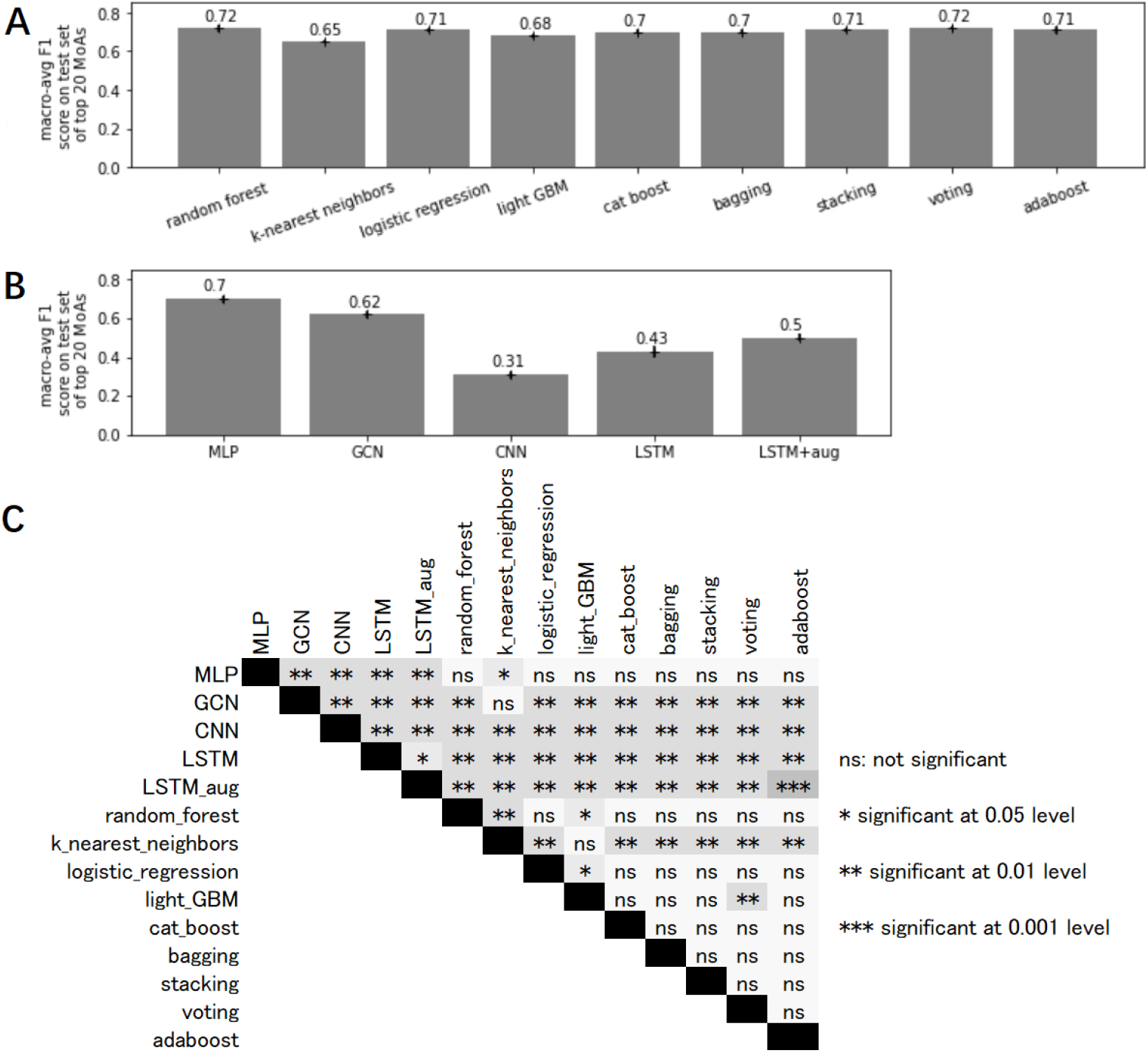
A). Comparison of macro-averaged F1 scores on the test set of the traditional machine learning models for the top 20 MoAs (i.e. the MoAs that were best represented in the data in terms of the number of compounds they had). B). Comparison of macro-averaged F1 scores on the test set of the deep learning models for the top 20 MoAs. C). Randomization test with Bonferroni correction of macro-averaged F1 scores on the test set of top 20 MoAs. The results are based on the averages across the nine shuffles of the training and validation data.

Test set F1 scores for the 10 selected MoAs (averaged across five shuffles of the training and validation data) comparing the MLP, trained on the compound structure data, EfficientNet, trained on the Cell Painting image data, and the global model, trained on both data sources are shown in Table 1. Our test set contained 24 compounds. This test set was the same for each of the shuffles of the training and validation data. For the MLP the F1 scores were very variable across the MoA classes, ranging from 0.08 for the JAK inhibitor test compounds to 1.00 for the Retinoid receptor agonist compounds. For EfficientNet the results were somewhat more stable, ranging from 0.48 for the Aurora kinase inhibitors to 0.98 for both the Protein synthesis inhibitors and the Retinoid receptor agonists. For the global model the results were even more stable, ranging from 0.68 for the ATPase inhibitors to 1.00 for the Retinoid receptor agonists. Our global model, achieving a macro-averaged F1 score of 0.92, revealed a clear additive/synergistic effect with an increase in F1 score of 0.11. The three different models were all significantly different from one another at the 5% significance-level based on randomization tests with Bonferroni corrected p-values. The predictive performance of the models for each of the test compounds is summarized in Table S1. We also show this comparison graphically in Figure 5 in which we have highlighted the compounds CBK290529 and CBK278556G, showing a very pronounced synergistic effect.

**Table 1.** F1 scores on the test set for the main three models explored for predicting the 10 selected MoAs. MLP used the chemical structure data, EfficientNet used the image data, and the Global model (see Figure 3) used both data formats. The results are based on the averages across the five shuffles of the training and validation data.

|  | MLP | EfficientNet | Global model |
| --- | --- | --- | --- |
| Aurora kinase inhibitor | 0.55 | 0.48 | 0.71 |
| Tubulin polymerization inhibitor | 0.37 | 0.92 | 0.97 |
| JAK inhibitor | 0.08 | 0.85 | 0.94 |
| Protein synthesis inhibitor | 0.54 | 0.98 | 0.99 |
| HDAC inhibitor | 0.98 | 0.89 | 0.99 |
| Topoisomerase inhibitor | 0.68 | 0.64 | 0.97 |
| PARP inhibitor | 0.51 | 0.96 | 0.98 |
| ATPase inhibitor | 0.64 | 0.58 | 0.68 |
| Retinoid receptor agonist | 1.00 | 0.98 | 1.00 |
| HSP inhibitor | 0.45 | 0.83 | 0.95 |
| Accuracy | 0.62 | 0.81 | 0.93 |
| Macro-averaged F1-score | 0.58 | 0.81 | 0.92 |
| Weighted-averaged F1-score | 0.61 | 0.81 | 0.93 |

**Figure 5.**
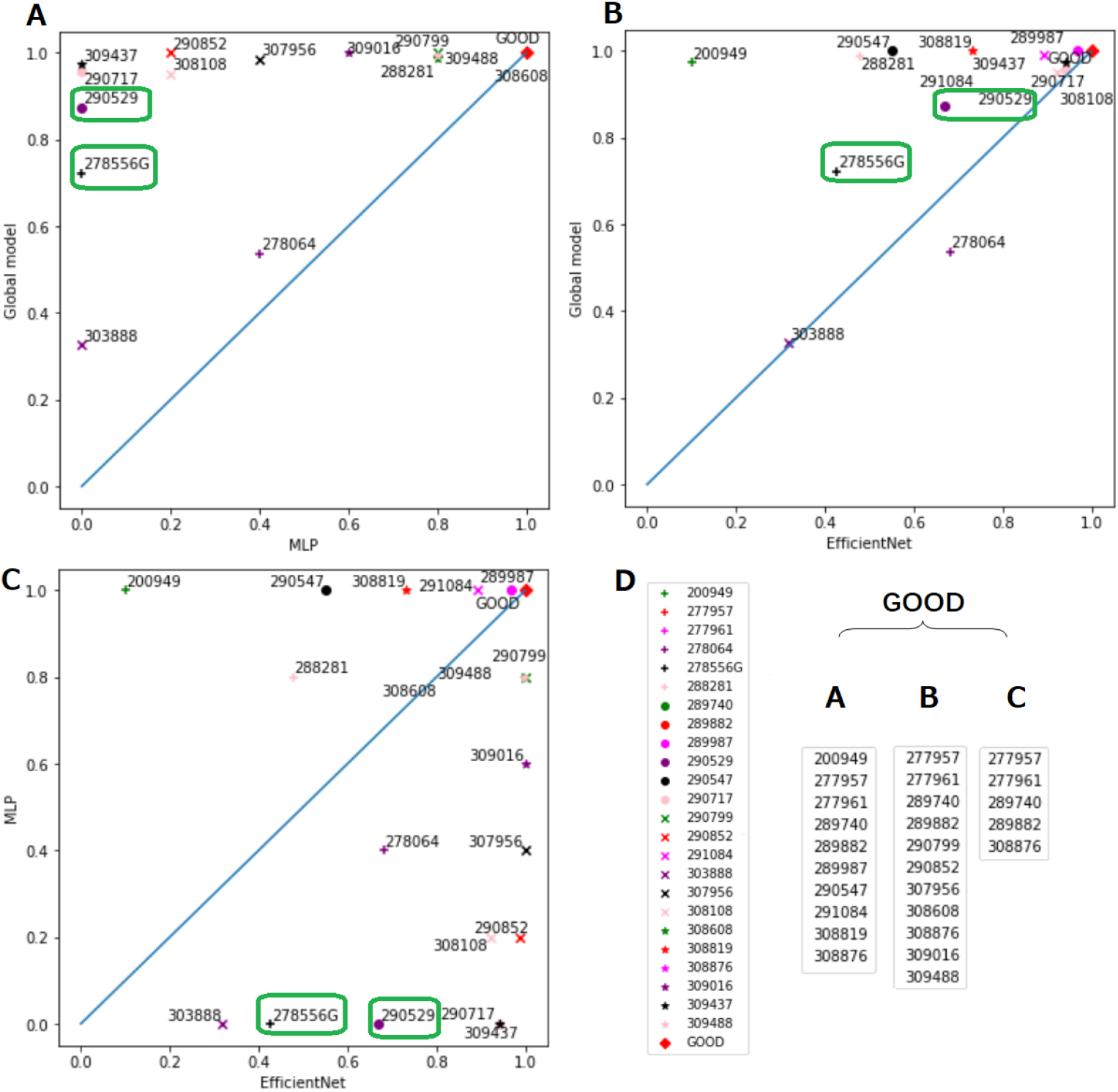
Comparison of the prediction rates for the three models for each compound: A). MLP, trained solely on the chemical structure data, versus the Global model, trained on both the chemical structure and Cell Painting image data; B). EfficientNet, trained solely on the image data, versus the Global model; C). EfficientNet versus MLP; D). ‘GOOD’ cluster contained compounds that possessed a prediction rate above 0.97 in panels A-C. Compounds CBK278556G and CBK290529 have been highlighted in green in panels A-C because they showed a greater synergistic effect than the other compounds.

## Discussion

We have introduced a novel and efficient approach for MoA prediction, which combines both Cell Painting image data and chemical structure data (in the form of Morgan fingerprints). Similarly to the study predicting cytotoxicity and proliferation [12] mentioned earlier we found that image based models outperformed those based on Morgan fingerprints and that integrating both data sources further boosted the performance. However, Lapins and Spjuth [10] found that chemical structure based models for MoA prediction were generally better than either L1000 or Cell Painting based models (note however that they used features derived from the images as opposed to the raw images themselves as model input). Chemical structure data (such as that obtained from Morgan fingerprints) can suffer from “Activity Cliffs”, whereby a small change in structure can result in a large difference in bio-activity, highlighting the need to supplement chemical structure data with additional sources of information [2], such as the Cell Painting image data used in the current study.

However, a few compounds were better predicted by the chemical structure based model, relative to the image based model. Perhaps the main disadvantage of image data such as Cell Painting is that not all compounds will necessarily produce a morphological change or the morphological effects may be very subtly and potentially masked by unaccounted for technical variations within and between plates during image capturing [2]. However, in the current study, to reduce potential bias caused by positional effects in the micro-well plates, the compounds and controls were distributed over the plates using PLAID and we standardized the images across the plates based on the control/DMSO wells. It is also possible that the compound does produce a morphological change but not in any of the cellular compartments or organelles captured using the Cell Painting assay. Another possibility is that the dose applied was not sufficient to produce a morphological change.

Concerning the traditional machine learning methods we explored for the chemical structure data, the ensemble methods outperformed the individual methods. Similarly, when combining a model based on multiple inputs, with separate modeling paths that come together to make a final prediction, we can potentially achieve better results than the models built on just one of the data categories. In this study, we showed this type of additive/synergistic effect by combining MLP for chemical structure data and EfficientNet for image data for MoA prediction.

Although it was somewhat surprising that for our models based on only the chemical descriptors, the simplest deep learning architecture, the MLP, outperformed the more complex networks architectures explored, a similar result has been obtained in a previous study [46] performing drug target prediction on a large benchmark dataset from the ChEMBL database. In our MLP architecture, we used an unconventionally high dropout rate to alleviate the overfitting problem as a result of the scarcity of chemical structure data. We also tested other possible architectures, such as reducing the dropout rate and increasing the number of hidden layers, with fewer neurons in each layer. However, these modifications did not improve the model performance.

Predicting the MoA of a compound can benefit greatly from the interaction of multiple sources of data [2]. Various studies [9,10,13,47] have shown that image and transcriptomics assays contain both overlapping and distinct cell state information. Thus, an even better predictive model than our final one could potentially be achieved with the additional integration of transcriptomics data. In the present study, for the image based model and the combined model, we used a set of ten well represented MoA classes. In future work we will explore the predictive ability of these models across a wider range of classes whilst accounting for potential polypharmacological effects. Contrary to previous belief, polypharmacology, where a compound concurrently engages with multiple targets or processes, is the rule rather than the exception in biology [19].

## Conclusion

In this work we used different models and data sources to predict MoA. The MLP was the best performing deep learning model for predicting MoA based on compound structures, and EfficientNet achieved a convincing result for predicting MoA based on Cell Painting image data. Integration of the MLP and EfficientNet, fitting to both datasets simultaneously, increased the F1 score by 0.11, thus exhibiting a clear additive/synergistic effect.

## Supporting Information

**Table S1.** Prediction rates for each compound in the test set for our three models. In the parenthesis first appears the compound ID, followed by the results of EfficientNet (trained on only the Cell Painting image data), the MLP (trained on only the compound structure data), and the Global model (trained on both datasets simultaneously).

|  |  |
| --- | --- |
| (CBK200949, 0.10, 1.00, 0.97) | (CBK277957, 1.00, 1.00, 1.00) |
| (CBK277961, 0.99, 1.00, 1.00) | (CBK278064, 0.68, 0.40, 0.54) |
| (CBK278556G, 0.43, 0.00, 0.72) | (CBK288281, 0.48, 0.80, 0.99) |
| (CBK289740, 0.99, 1.00, 1.00) | (CBK289882, 0.98, 1.00, 1.00) |
| (CBK289987, 0.97, 1.00, 1.00) | (CBK290529, 0.67, 0.00, 0.87) |
| (CBK290547, 0.55, 1.00, 1.00) | (CBK290717, 0.94, 0.00, 0.96) |
| (CBK290799, 1.00, 0.80, 1.00) | (CBK290852, 0.99, 0.20, 1.00) |
| (CBK291084, 0.89, 1.00, 0.99) | (CBK303888, 0.32, 0.00, 0.33) |
| (CBK307956, 1.00, 0.40, 0.99) | (CBK308108, 0.92, 0.20, 0.95) |
| (CBK308608, 1.00, 0.80, 0.99) | (CBK308819, 0.73, 1.00, 1.00) |
| (CBK308876, 0.99, 1.00, 1.00) | (CBK309016, 1.00, 0.60, 1.00) |
| (CBK309437, 0.94, 0.00, 0.97) | (CBK309488, 1.00, 0.80, 1.00) |

## Acknowledgments

We thank Jonne Rietdijk and Polina Georgiev for performing the Cell Painting assay and Anders Larsson for IT infrastructure assistance.

